# Anisotropic subdiffusion in the cytoplasm of living cells

**DOI:** 10.64898/2026.05.21.726834

**Authors:** Aranyak Sarkar, Pooja Yadav, Matthias Weiss

**Affiliations:** Experimental Physics I, University of Bayreuth, Universitätsstr. 30, D-95447 Bayreuth, Germany; Radiation and Photochemistry Division, Bhabha Atomic Research Centre, Mumbai 400085, India

## Abstract

The cytoplasm of eukaryotic cells is a complex fluid that harbors a variety of membrane-enveloped organelles with distinct spatial distributions and duties. Peroxisomes, for example, are vesicular entities with radii in the sub-micron range, dispersed all over the cytoplasm to provide oxidative reaction vessels across the cell. Here, we have tracked peroxisomes as biologically relevant tracer particles to explore the local structure of the cytoplasm. Focusing on the vast majority of (sub)diffusively moving peroxisomes, we show that these display a significant motion anisotropy even in the absence of cytoskeletal filaments or when the endoplasmic reticulum (ER) is disrupted. In untreated cells, the local preference direction aligns to a global axis that correlates with the cell’s long axis; disrupting the cytoskeleton or the ER progressively randomizes this alignment. Altogether, our data indicate that the cytoplasm features a nematic-like sub-structure rather than being an isotropic fluid.

---

The interior of eukaryotic cells is tidily organized into compartments (‘organelles’) that provide distinct chemical milieus for specific duties. Peroxisomes, for example, are membrane-enveloped organelles with diameters 0.1 − 1 *µ*m that host an oxidative environment in which macromolecules, e.g. long-chain fatty acids, are catabolized [1]. Peroxisomes are also key for regulating cellular hydrogen peroxide levels (hence their name), protecting the cell from uncontrolled/harmful oxidation events.

In mammalian cells, few tens to hundreds of peroxisomes are dispersed over the cytoplasm (cf. Fig. 1a for an example), most likely to support their role as omnipresent protective scavengers that eliminate toxic metabolic products. A dispersed state may also facilitate an equal partitioning of peroxisomes during cell division [2]. Key to this spatial arrangement are (transient) associations of peroxisomes with microtubules via suitable molecular motors that mediate long-range transport between cell center and the periphery (see [2] for a brief review). A recent study has shown, however, that only few percent of all peroxisomes show a ballistic motion along microtubules for more than five consecutive steps [3], whereas the vast majority rather shows a local and erratic motion in the cytoplasm.

**FIG. 1.**
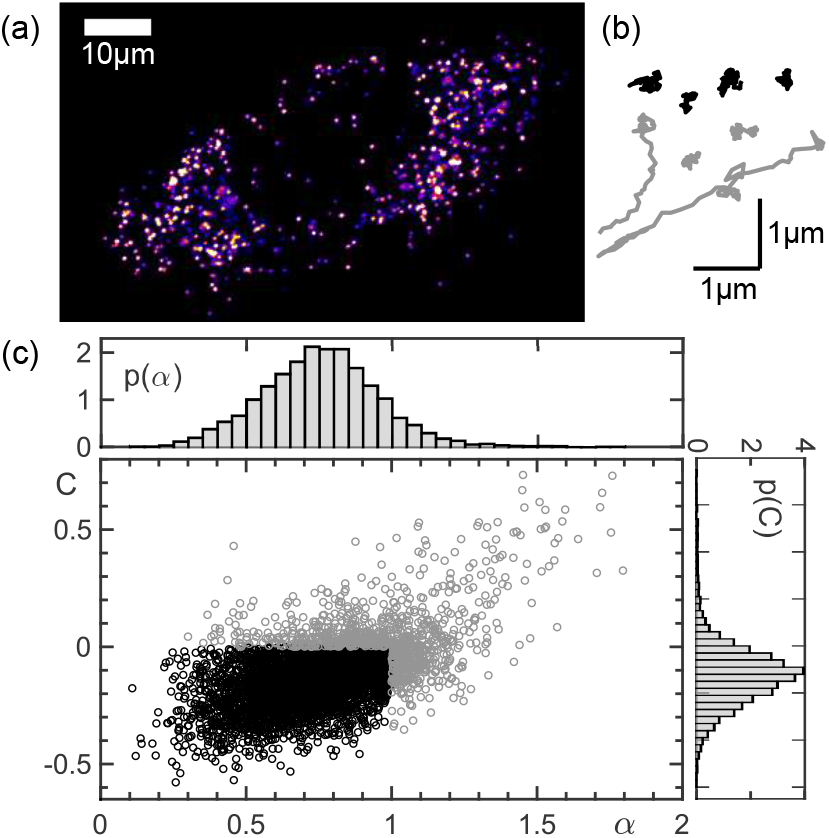
(a) Confocal fluorescence image of an U2OS cell with peroxisomes highlighted by SKL-GFP (movie in [8]). (b) Example trajectories of peroxisomes that were classified as (sub)diffusive and super-diffusive (black and grey, respectively). (c) Trajectories were classified as super-diffusive (grey) versus (sub)diffusive (black) when their TA-MSD scaling exponent was *α* > 1 or when the VACF value was *C* > 0.

Based on these observations, we hypothesized that peroxisomes could serve as prototypical test particles that report on local features of the cytoplasm as seen by a biologically relevant probe. Earlier work has shown that artificial tracer particles, which have been forced into cells, interact unspecifically with cytoplasmic structures, hence exhibiting a (partly intermittent) anomalous diffusion characteristic [4, 5]. Unlike artificial beads, peroxisomes are native probes that interact in a biologically regulated fashion with molecular motors, cytoskeletal filaments, and extended endomembrane structures like the endoplasmic reticulum [6] or mitochondria [7]). Therefore, one may expect that they can reveal biologically relevant features of the cytoplasm that remain hidden when monitoring artificial tracer particles.

Following this rationale, we have performed extensive single-particle tracking experiments on peroxisomes in living U2OS culture cells with a time resolution of Δ*t* = 110ms between successive frames (see Materials and Methods [8]). The in-plane motion of individual peroxisomes could be tracked over hundreds of frames (see Fig. 1b for examples) and trajectories with at least *N* = 100 consecutive positions (typically few thousands per cell) were used for subsequent analyses. In line with our previous approaches [5, 9], we first calculated for each trajectory the time-averaged mean square displacement (TA-MSD), ⟨*r*^2^(*τ*)⟩_*t*_ = ⟨(r(*t* + *τ*) − r(*t*))^2^⟩_*t*_, and their ensemble average (EA-TA-MSD) in a cell. All MSDs were fitted with a simple power law ⟨*r*^2^(*τ*) ⟩ = 4*Kτ*^*α*^ in the range *τ* ≤ 5 s [8]. Here, the scaling exponent *α* indicates (sub)diffusive (*α* ≤ 1) or super-diffusive (*α >* 1) motion, with *K* denoting the generalized diffusion coefficient. In addition, we calculated for each trajectory the characteristic value of the velocity autocorrelation function (VACF), *C* = ⟨v(*t*)v(*t* + Δ*t*)⟩_*t*_*/*⟨v(*t*)^2^⟩_*t*_, with the instantaneous velocity v(*t*) = [r(*t* + Δ*t*) − r(*t*)]*/*Δ*t*. While (sub)diffusion typically yields *C* ≤ 0, super-diffusive motion can be expected to yield *C >* 0 [9, 10].

In agreement with previous observations, we found that peroxisomes mostly show a (sub)diffusive motion, i.e. only 15-20% of all trajectories featured a super-diffusive signature (*α >* 1 or *C >* 0, see example in Fig. 1c). Dissecting trajectories into complementary sub-ensembles of (sub)diffusive (*α* ≤ 1 an *C* ≤ 0) and super-diffusive trajectories, the associated EA-TA-MSDs showed scaling exponents *α* ≈ 0.7 (similar to colloidal probes [4, 5]) and *α* ≈ 1.1, respectively. Due to the small super-diffusive pool, even the EA-TA-MSD without dissection showed a clear sublinear scaling (Fig. S1a in [8]).

Long-range transport of peroxisomes has been shown to depend on cytoskeletal motors [11]. Therefore, the small super-diffusive pool of trajectories can be rationalized to emerge by a simple stochastic switching between local diffusive motion and ballistic transport along cytoskeletal filaments. In contrast, the larger ensemble of (sub)diffusive trajectories, to which we will restrict our analysis in the remainder, can be expected to not include significant contributions of motor-driven transport along cytoskeletal elements.

In line with this notion, (sub)diffusive trajectories were seen to feature increments between successive frames, Δr = r(*t* + Δ*t*) − r(*t*), that appear consistent with an isotropic motion (see Fig. S1b in [8] for an example). However, transforming increments to polar coordinates, Δr → (Δ*r*_*ϕ*_, *ϕ*), and inspecting the angular probability distribution function (PDF), *p*(*ϕ*), regardless of the associated step length Δ*r*_*ϕ*_, a clear modulation with two distinct maxima of equal magnitude at *ϕ* = *ϕ*_0_ and *ϕ* = *ϕ*_0_ + *π* was seen in all cases (see example in Fig. 2a). This modulation indicates that increments of the (sub)difusive peroxisome trajectories have a directional preference with a nematic symmetry, i.e. bidirectional steps along an axis with orientation e_*ϕ*0_ are more probable than other directions. This observation for frame-to-frame steps not only persisted when analyzing steps taken within a period of 10Δ*t*, but the modulation was even stronger (Fig. 2a). To verify that the diffusion anisotropy is not caused by contaminations of few ballistic steps along the cytoskeleton (dispersed in mostly diffusive trajectories), we have also used a three-state hidden Markov model to complement our trajectory classification [8]. As a result, we observed that the anisotropy also persisted with this separation approach in all cases (Fig. S7 in [8]). Moreover, the anisotropy was seen to persist even when disrupting the cytoskeleton (see below).

**FIG. 2.**
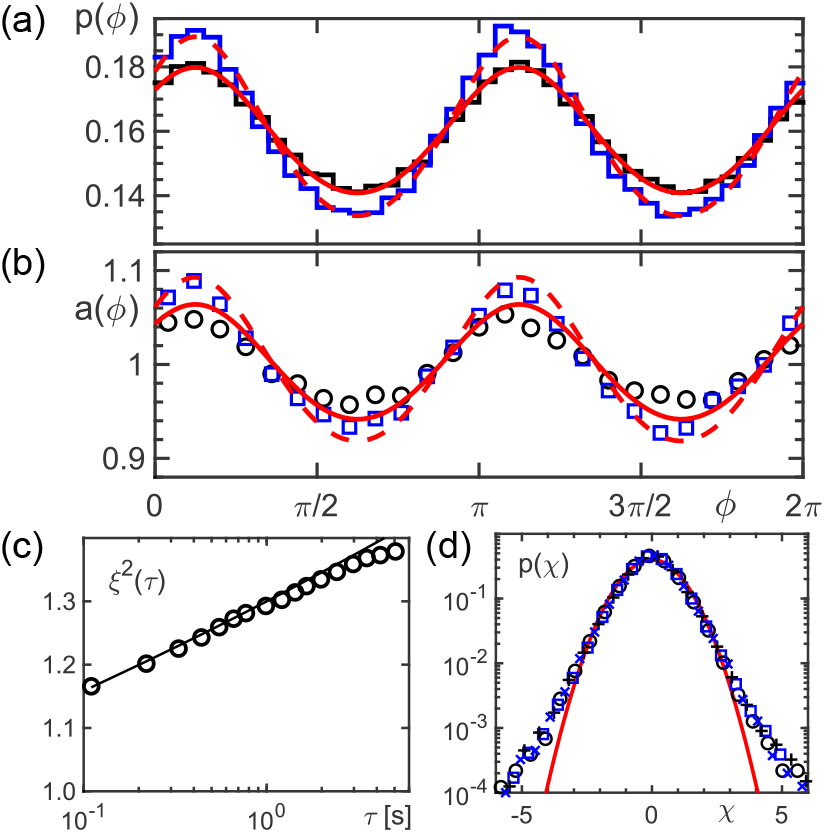
(a) The angular PDF *p*(*ϕ*) of steps taken by peroxisomes in the cell shown in Fig. 1a within one or ten frames (black and blue lines, respectively) features a marked modulation around the trivial value 1/(2*π*) with two distinct maxima of equal magnitude at *ϕ* = *ϕ*_0_, *ϕ*_0_ + *π* (*ϕ*_0_ ≈ *π*/8). This clear anisotropy of step directions in (sub)diffusive trajectories is well captured by Eq. (1) with *ξ* = 1.13 and *ξ* = 1.19 (full and dashed red lines), respectively. (b) Alongside, the relative step variation *a*(*ϕ*), defined in the main text, shows a similar modulation that is well captured by Eq. (2) without any additional parameter. (c) The squared step size anisotropy, *ξ*^2^(*τ*), defined as the ratio of EA-TA-MSDs ║ e_*ϕ0*_ and ⊥ e_*ϕ0*_, grows with the period *τ* in which the steps are taken (symbols; full line is an empirical fit *ξ*^2^(*τ*) = 1.3*τ* ^0.05^). (d) The PDF of normalized step increments, *p*(*χ*), is the same for steps ║ e_*ϕ0*_ and ⊥ e_*ϕ0*_ (black and blue symbols, respectively). Also the period in which the step was taken (crosses: *τ* = Δ*t*; open symbols: *τ* = 10Δ*t*) did not alter *p*(*χ*). In all cases, marked deviations from a normal distribution (red line) are seen, indicating a considerable heterogeneity of the (sub)diffusive motion within each trajectory.

As a complement to *p*(*ϕ*), we also analyzed the angle-resolved average step length *d*(*ϕ*) = ⟨Δ*r*_*ϕ*_⟩_*E*_. Normalizing *d*(*ϕ*) by its angle-averaged mean yields the relative step variation, *a*(*ϕ*) = *d*(*ϕ*)*/*⟨*d*(*ϕ*)⟩_*ϕ*_, that showed a very similar modulation as *p*(*ϕ*), again with equally high peaks at *ϕ*_0_ and *ϕ*_0_ + *π* (Fig. 2b). Therefore, peroxisome increments are not only more probable along the axis e_*ϕ0*_ but they are also longer. In accordance with our observations for *p*(*ϕ*), the modulation became stronger for steps taken within a period of 10Δ*t*. Thus, (sub)diffusive peroxisome trajectories clearly feature an anisotropic diffusion with larger steps along the axis e_*ϕ*0_.

To quantify the anisotropy increase for peroxisomal increments taken in periods *τ* = Δ*t*, 2Δ*t*, …, we have calculated the ratio 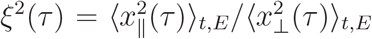 of one-dimensional EA-TA-MSDs along and perpendicular to the axis e_*ϕ0*_. In line with the data shown in Fig. 2a,b we observed a consistent growth of *ξ*^2^(*τ*) for increasing periods *τ* (Fig. 2c). This result further emphasizes that the (stochastic) process(es) that drive (sub)diffusive peroxisome motion are qualitatively and quantitatively different along and perpendicular to the axis e_*ϕ0*_.

Upon normalizing the steps Δ*x*_║_ and Δ*x*_⊥_ by the respective mean within single trajectories (enforcing mean step lengths ⟨*χ*_║_⟩ = ⟨*χ*_⊥_⟩ = 1 for each trajectory), information on the trajectories’ spatiotemporal heterogeneity can be obtained [9]. As a result, we observed that the shape of all PDFs *p*(*χ*) neither depended on the direction (║ or ⊥ e_*ϕ0*_) nor on the period *τ* in which the step was taken (Fig. 2d). However, all PDFs strongly deviated from a normal distribution, indicating that (sub)diffusive peroxisome motion features a considerable spatiotemporal heterogeneity in all directions.

To obtain a meaningful interpretation of our experimental data, we have formulated a simple model (see also sketch in Fig. 3a): We assume a Gaussian PDF of step increments along and perpendicular to the preferred direction e_*ϕ0*_, Δ*x*_║_ and Δ*x*_⊥_, with variances 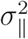 and 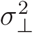, such that the squared anisotropy 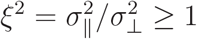 1 is consistent with our above defintion of *ξ*^2^(*τ*). Transforming Euclidean to polar coordinates yields the PDF *p*(*ρ, ϕ*) for radial step increments of length *ρ* along the direction *ϕ*. From this, the PDF of steps of any length in direction *ϕ* can be calculated as

**FIG. 3.**
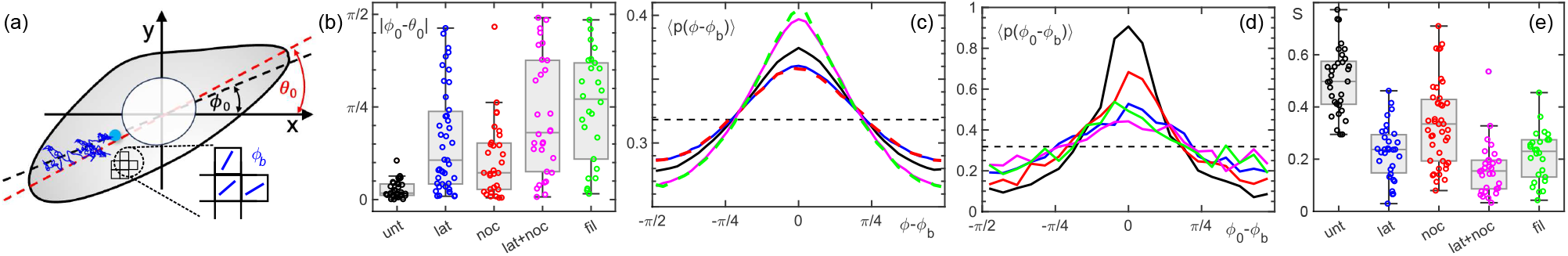
(a) Sketch of the orientations of trajectories and the cell’s long axis. Squares indicate the bins for defining the local directors *ϕ*_*b*_ [Eq. (3)]. (b) The angle difference |*ϕ*_0_ − *θ*_0_| between the preferential direction of peroxisome motion, e_*ϕ0*_, and the long cell axis, e_*θ0*_, is on average only 5° in untreated cells (‘unt’, black symbols). For latrunculin- and nocodazole-treated cells (‘lat’ and ‘noc’, blue and red symbols) the deviations are considerably larger but deviate on average only by about 30°. Upon treatment with latrunculin and nocodazole (‘lat+noc’, magenta) or with filipin (‘fil’, green), both of which disrupt the ER, the deviations grow to more than 45°, indicating a strong randomization of the axes’ relative orientation. (c) Cell-ensemble averages of *p*(*ϕ* − *ϕ*_*b*_) feature a significant modulation and hence a local anisotropic diffusion for all conditions (same color code as before). Dashed line: expectation for isotropic motion. (d) The ensemble-averaged PDF ⟨*p*(*ϕ*_0_ − *ϕ*_*b*_)⟩ indicates a strong alignment of local directors *ϕ*_*b*_ to a global director *ϕ*_0_ for untreated cells. Although softened when disrupting the cytoskeleton or the ER, a marked modulation remains. (e) The nematic order parameter *S* quantifies the latter, indicating a substantial alignment of *ϕ*_*b*_ and *ϕ*_0_ for all conditions.

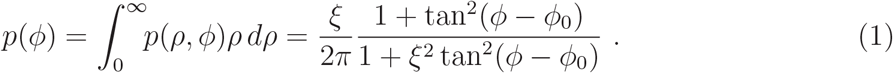

This expression fits our experimental data very well (Fig. 2a), when adjusting the value of *ξ* to the period *τ* between successive steps. In addition, one can calculate from Eq. (1) the angular variation of the mean step length via the first moment of *p*(*ρ, ϕ*),

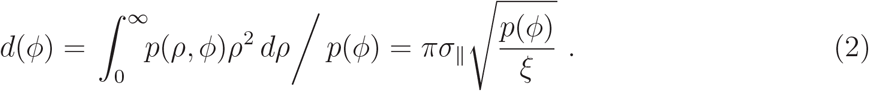

Dividing *d*(*ϕ*) by its angular average (*d*(*ϕ*))_*ϕ*_ yields, as before, *a*(*ϕ*). The resulting analytical expression fits our experimental data on *a*(*ϕ*) very well (Fig. 2b), without any additional free parameter. To rationalize the experimentally seen growth of 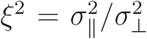 (cf. Fig. 2c) one may consider an isotropic and anti-persistent fractional Brownian motion (FBM) process for Δ*x*_║_ and Δ*x*_⊥_, onto which a one-dimensional Markovian random walk along the direction e_*ϕ0*_ is superimposed. While the FBM induces an isotropic subdiffusive motion with 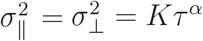 and *α <* 1, the additional Markovian steps along e_*ϕ0*_ not only elevate the diffusion coefficient *K* for 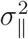 but they also perturb the FBM’s anti-persistent memory. Consequently, a (slightly) higher value of the scaling exponent *α* along the preferred direction e_*ϕ0*_ emerges, leading to a time-dependent increase of *ξ*^2^.

The observation that each cell features a preferential direction e_*ϕ0*_ prompted us to explore the underlying origin. Visual inspection suggested that the long axis of each cell’s gyration ellipsoid, e_*θ*0_ (obtained from the gyration tensor of the convex hull of all peroxisome positions), may correlate with e_*ϕ0*_; (cf. sketch in Fig. 3a). Extracting the angle difference 0 ≤ |*ϕ*_0_ − *θ*_0_| ≤ *π/*2 between e_*ϕ0*_ and e_*θ0*_, we indeed observed only minor deviations between these two axes, in the range of few degrees (Fig. 3b). Thus, the global orientation of individual cells appears to determine the preferential direction of (sub)diffusive transport of peroxisomes. This observation suggests that extended structures like cytoskeletal filaments or the endoplasmic reticulum network (ER), are involved in setting a glocal director field *ϕ*_0_ for the observed (sub)diffusion anisotropy.

Before testing this hypothesis by disrupting these structures with specific drugs, we have probed the intracellular variation of the (sub)diffusion anisotropy by analyzing its local director field. To this end, we calculated for all (sub)diffusive trajectories within a cell all frame-to-frame increment vectors in polar coordinates, r(*t* + Δ*t*) − r(*t*) → (Δ*r, ϕ*). The resulting angles *ϕ* were assigned to the respective average positions r^⋆^(*t*; *τ*) = [r(*t* + *τ*) + r(*t*)]*/*2. Partitioning the plane into square bins of edge length 1 *µ*m, the positions r^⋆^(*t*; *τ*) were discretized and data for *ϕ* were assigned to the respective bins, yielding *N*_*b*_ angles in bin *b*. To exclude statistically poor bins from subsequent analyses, we only considered bins with *N*_*b*_ ≥ 5. With this, we defined a local director field *ϕ*_*b*_ that determines the locally preferred direction in bin *b* (cf. Fig. 3a):

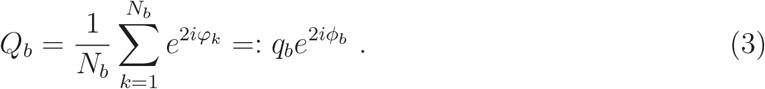

Given that we are dealing with a nematic order, angle shifts *ϕ* → *ϕ* + *π* must not change the order parameter, hence the factor 2 in the exponent. An example for the local director field *ϕ*_*b*_ is shown in Fig. S1c in [8], supporting the impression that the local variation of *ϕ*_*b*_ is very low. To quantify the robustness of our findings, beyond the single cell example, we analyzed an ensemble of *n* = 33 untreated cells with *m* = 52818 trajectories. As a result, we observed that (sub)diffusive step increments in all cells showed a clear anisotropy, highly similar to Fig. 2. In particular, the (sub)diffusive steps in all cells nicely followed a local director *ϕ*_*b*_, indicated by a marked modulation in the PDF *p*(*ϕ* − *ϕ*_*b*_) (Fig. 3c). In addition, the local directors *ϕ*_*b*_ closely followed the cell-specific global director, *ϕ*_0_, as evidenced by the ensemble-average PDF (*p*(*ϕ*_0_ − *ϕ*_*b*_)) (Fig. 3d) and by the corresponding nematic order parameter *S* = (cos(2[*ϕ*_*b*_−*ϕ*_0_])) that assumed values around *S* = 0.5 (Fig. 3e). Together with the correlation *ϕ*_0_ ≈ *θ*_0_ (Fig. 3b), this confirms an anisotropic (sub)diffusion of peroxisomes also for an ensemble of cells. As in individual cells, the anisotropy has the same local and global preference axis that correlates with the long axis of the cell. Separating (sub)diffusive step increments with a three-state hidden Markov model, as an alternative to the selection via *α* and *C*, yielded highly similar results [8]. We therefore conclude that the observed anisotropy is very similar at all cell loci, suggesting that the diffusion anisotropy emerges from local structures that adapt a cell-wide directional preference.

Next, we inspected ensembles of cells in which cytoskeletal filaments or the ER were disrupted by specific drugs [8]. Analyses of single example cells (corresponding to Fig. 1 and Fig. 2) are summarized in Figs. S2-S5 [8]. Given that actin filaments are known to be key for cell adhesion and shape, even aligning with the major cell axis (see, e.g., [12]), we anticipated changes in the (sub)diffusion anisotropy when disrupting actin filaments by the established drug latrunculin A [13] (*n* = 50 cells, *m* = 33082 trajectories). Somewhat unexpectedly, however, the modulation in *p*(*ϕ* − *ϕ*_*b*_) was only softened slightly (Fig. 3c), indicating again a strong anisotropic (sub)diffusion with local directors *ϕ*_*b*_. As compared to untreated cells, however, the alignment of the local directors to a global direction was less strong, as evidenced by the broadened PDF (*p*(*ϕ*_0_ − *ϕ*_*b*_)) and a reduced nematic order parameter *S* (Fig. 3d,e). In addition, the correlation of e_*ϕ0*_ and e_*θ0*_ was less strong (Fig. 3b). Treatment with latrunculin A was also seen to slightly reduce the fraction of trajectories with a super-diffusive signature and to alter the PDF of step lengths between successive frames, *p*(Δ*r*), for (sub)diffusive trajectories, featuring markedly larger values and an increased mean as compared to untreated cells (Fig. S6 [8]). Thus, our data provide evidence that actin filaments are not the dominant cause of the observed anisotropic (sub)diffusion of peroxisomes. Rather, intact actin filaments appear to hamper the motion of peroxisomes.

We then applied the drug nocodazole via an established protocol [14, 15] to disrupt all microtubules (*n* = 31 cells, *m* = 67471 trajectories). Microtubules are best known as filaments along which organelles are transported. In line with this notion, we observed that almost no trajectories with a super-diffusive signature remained (with values for *α* and *C* near to the selection threshold). Still, we observed a similar modulation in *p*(*ϕ* − *ϕ*_*b*_) as for latrunculin-treated cells (Fig. 3c) and an even better alignment of the local directors to a global direction (Fig. 3d,e). Therefore, even in the absence of microtubules, the modulation remained significant and also the correlation between e_*ϕ0*_ and e_*θ0*_ remained similar to latrunculin-treated cells (Fig. 3b). The PDF of (sub)diffusive steps, *p*(Δ*r*), hardly deviated from that of untreated cells (Fig. S6 [8]). From these data, we conclude that microtubules might contribute to the observed anisotropy, but they are not the sole or even dominant cause for its emergence.

Combined treatment with nocodazole and latrunculin A (*n* = 30 cells, *m* = 25996 trajectories), which cells tolerate only for a very limited time before dying off, resulted in a further dealignment of local and global directors and a further randomization between *ϕ*_0_ and *θ*_0_ (Fig. 3). On local scales, however, the (sub)diffusion anisotropy persisted and the modulation in *p*(*ϕ* − *ϕ*_*b*_) was even slightly enhanced (Fig. 3). Therefore, the cytoskeleton cannot be held responsible for the locally occuring anisotropy, but the data suggests, at first glance, that the combined cytoskeleton is the major organizer for aligning local directors to each other. Yet, we also observed that the ER was severely compromised in these cells.

Hypothesizing that the ER is the important structure, rather than the filaments, we selectively disrupted the ER network by applying filipin [16]. The result (*n* = 26 cells, *m* = 7405 trajectories) was very similar to the combined treatment with nocodazole and latrunculin A (Fig. 3), providing strong evidence that the ER, rather than the cytoskeleton, is the major organizer for aligning the local directors *ϕ*_*b*_ to a global director *ϕ*_0_. Again, the modulation in *p*(*ϕ* − *ϕ*_*b*_) was higher than in untreated cells, indicating a substantial (sub)diffusion anisotropy.

We speculate that this enhancement is due to the fragmentation of the ER into small membrane structures that are scattered across the cytoplasm, complementing already existing membrane structures, such as mitochondria or endosomes. All of these interfaces may locally block and guide movements of neighboring particles, hence enforcing a local anisotropy for diffusional motion of neighboring particles, akin to the properties of a nematic fluid. Without a cell-spanning structure, these constraints remain local and do not get aligned, whereas an intact ER may organize local director fields to a global preference direction. The situation is hence reminiscent to the initial magnetization of a ferromagnetic material: spontaneously formed ferromagnetic domains (here: nematic regions with director *ϕ*_*b*_) need to be aligned by an external field (here: the ER) to form a preferred direction on macroscopic scales (here: *ϕ*_0_). In any case, cytoskeletal filaments are dispensable for the local anisotropy and also appear unessential for the emergence of a global direction for anisotropic (sub)diffusion. Based on these considerations, we expect that not only peroxisomes but virtually all particle-like structures in the cytoplasm will exhibit a similar anisotropy signature, organized by the ER network.

## ACKNOWLEDGMENTS

The authors thank D. Grätz and K. Weidner-Hertrampf for technical support, and Fred Wouters (University of Goettingen) for sharing the plasmid for SKL-GFP. Financial support by the VolkswagenStiftung (Az. 92738) and by the Elite Network of Bavaria (Study Program Biological Physics) are gratefully acknowledged.

