## supplemental text & figures for "Anisotropic subdiffusion in the cytoplasm of living cells"

Aranyak Sarkar

*Experimental Physics I, University of Bayreuth, Universitätsstr. 30, D-95447 Bayreuth, Germany and Radiation and Photochemistry Division, Bhabha Atomic Research Centre, Mumbai 400085, India*

Pooja Yadav and Matthias Weiss*

*Experimental Physics I, University of Bayreuth, Universitätsstr. 30, D-95447 Bayreuth, Germany*

### I. MATERIALS AND METHODS

#### A. Cell culture and microscopy

Bone osteosarcoma cells (U2OS, DSMZ ACC-832, RRID: CVCL 0042) were cultured as described before (Speckner *et al.*, 2024). In brief, cells were grown in T-25 flasks (BioLite No. 130189 Thermo Scientific, Germany) at 5% $CO_2$ and $37°C$, using Dulbecco's Modified Eagle's Medium (DMEM, Sigma-Aldrich D5671) supplemented with 10% fetal bovine serum (Sigma-Aldrich F7524), 5% sodium pyruvate solution (Sigma-Aldrich S8636) and 5% penicillin/streptomycin (Sigma-Aldrich $P4333$). Cells were split at 80% confluency every two to three days using pre-warmed trypsin EDTA (0.25%, Sigma-Aldrich T4049) for 3 min to detach cells.
For microscopy, U2OS cells were seeded at a density of 15,000 cells per well in 2-well plate $\mu$-Dishes (ibiTreat, 80286). To visualize peroxisomes, cells were transfected with a GFP plasmid carrying an additional serine-lysine-leucine (SKL) motif, targeting the protein to peroxisomes (SKL-GFP, a kind gift from Fred Wouters, University of Göttingen). Transfection was performed 24 h prior to microscopy using FuGENE transfection reagent for 15 min in 100 $\mu$L serum-free Opti-MEM (Gibco, Germany, 31985062), and 100 $\mu$L of the transfection solution was added dropwise to each well. Prior to imaging, cells were washed twice in Dulbecco's phosphate-buffered saline (Gibco, Germany, 14190144) and then supplemented with $CO_2$-independent imaging medium (MEM without phenol red, supplemented with 5% HEPES and 1% penicillin/streptomycin).
To depolymerize microtubules, we used a previously described and validated protocol (Hirschberg *et al.*, 1998; Stadler and Weiss, 2017). In brief, the imaging medium was supplemented with 10 $\mu$M nocodazole (Sigma-Aldrich, Germany, M1404), which blocks microtubule growth. For this, 2 $\mu$M stock solution of nocodazole (Sigma Aldrich, M1404), dissolved in dimethylsulfoxide (DMSO, AppliChem, Germany, A3672), was diluted to a working concentration of 10 $\mu$M. Cells were chilled on ice for 10 min to induce microtubule depolymerisation before incubating for another 24 h at 37°C with 5% $CO_2$. Imaging was also performed in the presence of nocodazole to prevent a *de novo* growth of microtubules. To disrupt actin filaments, cell and imaging media were supplemented with latrunculin A (TOCRIS Bioscience, CAS No. 76343-93-6; dissolved in DMSO) in a final concentration of 5 $\mu$M (2 h prior to imaging). To depolymerize both, microtubules and actin filaments, cells were treated with 10 $\mu$M nocodazole for 4h prior to imaging, following the protocol mentioned above. Latrunculin A (concentration 5 $\mu$M) was added two hours after the application of nocodazole, and cells were incubated for another two hours prior to imaging. Imaging was performed in the presence of both drugs. To disrupt the endoplasmic reticulum, filipin III (Sigma-Aldrich,F4767) was used at a working concentration of 15 $\mu$g/ml to disrupt the endoplasmic reticulum (Axelsson and Warren, 2004). Prior to the treatment, cells were washed twice with Dulbecco's phosphate-buffered saline and then exposed to filipin III for 30 min at room temperature and then incubated for another 30 min at 37°C before imaging.
Prior to imaging, cells were washed twice in Dulbecco's phosphate-buffered saline (# 14190144 Gibco, Germany) and then supplemented with imaging medium. Imaging was performed at 37°C with a customized spinning-disk confocal microscope, consisting of a Leica DMI 4000 microscope body (Leica Microsystems, Germany), a CSU-X1 (Yokogawa, Japan) spinning disk unit, and a custom-made incubation chamber. Images were acquired using an Evolve 512 EMCCD camera (Photometrics, USA)paired with an HC PL APO 63x/1.4 (Leica Microsystems) oil immersion objective, resulting in a pixel size of 56.2 nm. Samples were illuminated at 488 nm and fluorescence light was detected in the range 500-550 nm. The whole setup was controlled by custom written LabView software (National Instruments, USA). For single-particle tracking of peroxisomes, time-resolved image stacks were taken at an interval of $\Delta t = 110$ ms using a $2 \times 2$ pixel binning. Representative movies for peroxisomes in untreated and nocodazole-treated cells are provided as supplement.

---

*Contact: ``

#### B. Image and trajectory analysis

Bright GFP-tagged peroxisomal puncta were localized in each frame of the raw image stack using *THUNDERSTORM* in Fiji/ImageJ in a two-stage workflow. Here, candidate emitters were first detected after B-spline wavelet filtering (spline order 4, scale 3) to yield pixel-precision seeds, and then refined to sub-pixel accuracy by maximum-likelihood fitting of an integrated 2D Gaussian point-spread function with a 2-pixel fitting radius, returning $(x, y)$ positions in each frame with subdiffraction precision. The localization tables were exported to MATLAB for frame-to-frame association via a nearest-neighbor linker that restricted links to consecutive frames without gap closing (any missed detection terminated the track and any subsequent reappearance initiated a new trajectory). To this end, a maximum per-frame displacement of 1 $\mu$m was used (setting an upper-speed bound of $\sim 10$ $\mu$m/s, well above the typical peroxisome speed ($\lesssim 1$ $\mu$m/s). Trajectories were only retained for subsequent analyses when they consisted of at least 100 consecutive frames. Our approach was seen (in comparison, for example, to FIJI/Trackmate) to minimize localization errors and spurious assignments, to eliminate gap-closing ambiguities, and to ensured adequate temporal depth for robust downstream analyses. Whenever a peroxisome was lost, e.g. when moving out of focus, the respective trajectory was closed, i.e. lost positions were not interpolated. In total, we analyzed $n = 33$ (untreated), $n = 31$ (nocodazole-treated), and $n = 50$ (latrunculin-treated) cells that yielded a total of $m = 52818$, $m = 67471$, and $m = 33082$ trajectories with at least $N = 100$ consecutive positions.
Particle trajectories were analyzed in Matlab (Matlab 2018b, The MathWorks Inc., USA) using newly developed scripts as well as our previously introduced toolbox of analysis routines (Rehfeldt and Weiss, 2023). TA-MSDs were fitted with a simple power law in the range $\Delta t \leq \tau \leq 5$ s, as stated in the main text, using an equidistant spacing of lag times on a logarithmic scale to compensate the successive growth of available MSD values for increasing lag times.

#### C. Additional notes on figures

Scale bars in all microscopy images (Fig. 1a and Figs. S2-5a) denote 10 $\mu$m. Swarm plots of data in Fig. 3b and Fig. 3e of the main text were produced with the Matlab routine *swarmchart*. Here, data points are jittered based on their kernel density estimate (similar to violin plots). Superimposed box plots were produced with the Matlab routine *boxchart*. Here, the box denotes the 25% and 75% quantile of the data, with the median shown as horizontal line within the box. Data points are classified as outliers, when they are not within a distance of 1.5× the interquartile range from the minimum/maximum of the box. Whiskers extend to the minimum and maximum data values that are not classified as outliers.

### II. SEPARATION OF MOBILITY STATES BY A THREE-STATE HIDDEN MARKOV MODEL

To complement the separation of motion states via the TA-MSD scaling and the VACF, we have also used a three-state hidden Markov model (HMM) as an alternative. We start by defining any displacement vector within a period $L\Delta t$ in Euklidean and polar coordinates,

$$\Delta \mathbf{r}_n = \mathbf{r}_{n+L} - \mathbf{r}_n = (\Delta x_n, \Delta y_n) = (\rho_n, \phi_n) \ . \tag{1}$$

From the set of displacement vectors obtained for (sub)diffusively rated trajectories within a cell, we formally define the cell's global nematic reference orientation via

$$\mu_g = \frac{1}{2} \arg \left( \left\langle e^{i2\phi_n} \right\rangle \right) \ . \tag{2}$$

To account for spatial heterogeneity, the cell is then partitioned into spatial bins $B_{ij}$ (1 $\mu$m$^2$ squares) for which local director fields are constructed. To this end, each displacement $\Delta \mathbf{r}_n$ is assigned to the midpoint of the involved positions, $\mathbf{x}_n = [\mathbf{r}_{n+L} + \mathbf{r}_n]/2$, which uniquely belongs to one of the bins. With this, each bin contains $N_{ij}$ displacement vectors from which we compute the local director field $\mu_{ij} = \frac{1}{2} \arg(Q_{ij})$ with $Q_{ij} = \sum_{n=1}^{N_{ij}} e^{i2\phi_n}$; for bins with a poor statistics, i.e. $N_{ij} < N_{\min} = 10$, we set $\mu_{ij} = \mu_g$. With this, every displacement vector $\Delta \mathbf{r}_n$ is associated with the local director field $\mu_{ij}$ in its bin $B_{ij}$ and the displacement can be expressed in this local coordinate system:

$$\Delta r_{\parallel,n} = \Delta x_n \cos \mu_{ij} + \Delta y_n \sin \mu_{ij} \quad \Delta r_{\perp,n} = -\Delta x_n \sin \mu_{ij} + \Delta y_n \cos \mu_{ij}. \tag{3}$$

All subsequent state inferences are performed in this local frame.
In total we consider three distinct states that particle displacements belong to. Denoting the hidden state as $z_n \in \{1, 2, 3\}$, the corresponding observation ('emission') is $\mathbf{o}_n = (\Delta r_{\parallel,n}, \Delta r_{\perp,n})$.

**State 1** describes rest or tethered motion, when peroxisomes are bound to microtubules but do not move. The output probability is hence an anisotropic zero-centered Gaussian,

$$b_1(\mathbf{o}_n) = \mathcal{N}(\Delta r_{\parallel,n}; 0, s_{1\parallel}^2)\,\mathcal{N}(\Delta r_{\perp,n}; 0, s_{1\perp}^2), \qquad s_{1\parallel} \geq s_{1\perp} \ . \tag{4}$$

Here and below, $\mathcal{N}(x; m, s^2)$ denotes a Gaussian PDF for the observable $x$ with a mean $m$ and variance $s^2$.
**State 2** describes isotropic passive diffusion,

$$b_2(\mathbf{o}_n) = \mathcal{N}(\Delta r_{\parallel,n}; 0, s_2^2)\,\mathcal{N}(\Delta r_{\perp,n}; 0, s_2^2). \tag{5}$$

**State 3** finally describes directional runs along the local nematic axis, modeling that peroxisomes are transported along microtubules. To preserve nematic, rather than polar, symmetry, we use here a symmetric two-component mixture,

$$b_3(\mathbf{o}_n) = \left[\frac{1}{2}\mathcal{N}(\Delta r_{\parallel,n}; +u_3, s_{3\parallel}^2) + \frac{1}{2}\mathcal{N}(\Delta r_{\parallel,n}; -u_3, s_{3\parallel}^2)\right]\mathcal{N}(\Delta r_{\perp,n}; 0, s_{3\perp}^2), \tag{6}$$

where $u_3 > 0$ sets the run velocity along the local axis.
The emission model is parameterized by

$$\Theta_{\text{em}} = \{w_1, w_2, w_3,\ s_{1\parallel}, s_{1\perp},\ s_2,\ u_3, s_{3\parallel}, s_{3\perp}\} \ , \tag{7}$$

with $w_k$ the mixture weights for the three states. These parameters are first estimated from the pooled projected steps by expectation-maximization, providing a stable initialization for the HMM while keeping the state identities fixed across trajectories.
Temporal structure is then introduced through a three-state first-order Markov chain,

$$A_{ij} = P(z_{n+1} = j \mid z_n = i), \qquad \pi_i = P(z_1 = i), \qquad i, j \in \{1, 2, 3\}. \tag{8}$$

For a trajectory of $T$ lagged steps, the complete-data likelihood is

$$P(z_{1:T}, \mathbf{o}_{1:T}) = \pi_{z_1} b_{z_1}(\mathbf{o}_1) \prod_{n=2}^{T} A_{z_{n-1}z_n}\, b_{z_n}(\mathbf{o}_n). \tag{9}$$

Given fixed emissions, the transition matrix $A$ and initial distribution $\pi$ are estimated by the Baum–Welch algorithm. With

$$\gamma_n(i) = P(z_n = i \mid \mathbf{o}_{1:T}), \qquad \xi_n(i, j) = P(z_n = i, z_{n+1} = j \mid \mathbf{o}_{1:T}), \tag{10}$$

the transition update is

$$A_{ij} = \frac{\sum_{n=1}^{T-1} \xi_n(i, j)}{\sum_{n=1}^{T-1} \gamma_n(i)}. \tag{11}$$

The most likely hidden-state sequence is then obtained by Viterbi decoding.
As a result, this construction yields a state-space decomposition in the local nematic frame: state 1 captures anisotropic zero-mean fluctuations, state 2 isotropic passive motion, and state 3 bidirectional run-like transport along the local axis. These states hence reflect the simplest decomposition of peroxisome motion into ballistic motion along the cytoskeleton (state 3), pausing while being bound to the cytoskeleton (state 1), and a passive isotropic diffusion while not being bound to the cytoskeleton (see examples in Fig. S7). Please note that we deliberately chose a bidirectional emission for the run state, although many runs may be locally unidirectional. This conservative choice effectively over-represents run-like motion, biasing the model to absorb ambiguous aligned steps into the run state rather than into the passive or subdiffusive states. Consequently, any motion retained in the passive/diffusive class cannot be attributed to an overly restrictive definition of runs.
Application of this three-state HMM scheme resulted in the finding that even those displacement vectors that were most consistent with the simple isotropic diffusion state (state 2) showed the same anisotropy signature as the other two states (see Fig. S7). These data are in line with our separation of trajectories into a (sub)diffusive pool and its complement via the TA-MSD and the VACF. We therefore conclude that all displacement vectors, including the passive ones, have the same anisotropy.

### III. CAPTIONS FOR SUPPLEMENTAL MOVIES

**untreated_trimmed_2min.mp4**
Time-lapse movie of peroxisome motion in the untreated cell shown in Fig. 1, trimmed to 2 min length.

**lattreated_trimmed_2min.mp4**
Time-lapse movie of peroxisome motion in the latrunculin-treated cells shown in Fig. S2, trimmed to 2 min length.

**noctreated_trimmed_2min.mp4**
Time-lapse movie of peroxisome motion in the nocodazole-treated cells shown in Fig. S3, trimmed to 2 min length.
**noclattreated_trimmed_2min.mp4**
Time-lapse movie of peroxisome motion in the nocodazole-latrunculin-treated cells shown in Fig. S4, trimmed to 2 min length.

**filtreated_trimmed_2min.mp4**
Time-lapse movie of peroxisome motion in the filipin-treated cells shown in Fig. S5, trimmed to 2 min length.

### IV. SUPPLEMENTAL FIGURES

#### Fig. S1

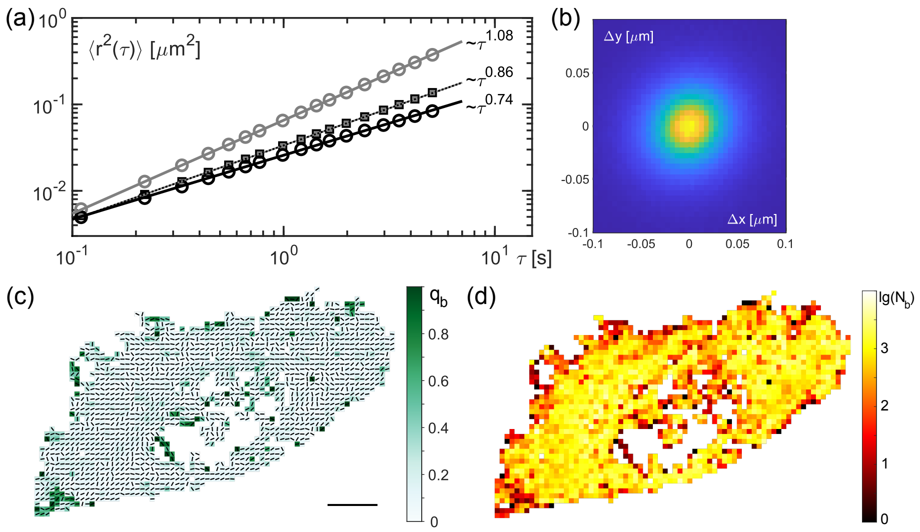


FIG. S1 Additional data for the cell shown in Fig. 1a of the main text. (a) The EA-TA-MSDs of (sub)diffusive and superdiffusive trajectory sub-ensembles (black and grey circles) displays a distinct sub- and superlinear scaling, respectively. The unseparated ensemble of trajectories still features a sublinear scaling (small squares). (b) The associated density plot of increment vectors, $\Delta\mathbf{r} = \mathbf{r}(t+\Delta t) - \mathbf{r}(t) = (\Delta x, \Delta y)^T$, does not indicate a major deformation from which one may conclude an anisotropic motion of peroxisomes. (c) Field of local directors $\phi_b$ (obtained from $Q_b$ via Eq. (3), main text) shown as black quivers. Superimposed (color-coded) is the associated amplitude $q_b$, featuring a median value of 0.1. (d) Color-coded local number $N_b$ of phases $\varphi$ that entered $Q_b$ (please note the log-scale). The median number of phases per bin was 694, yielding an estimate that even the small values for $q_b$ are above the null result for random/scrambled phases (i.e. $q_b > 1/\sqrt{694}$).

**Fig. S2**

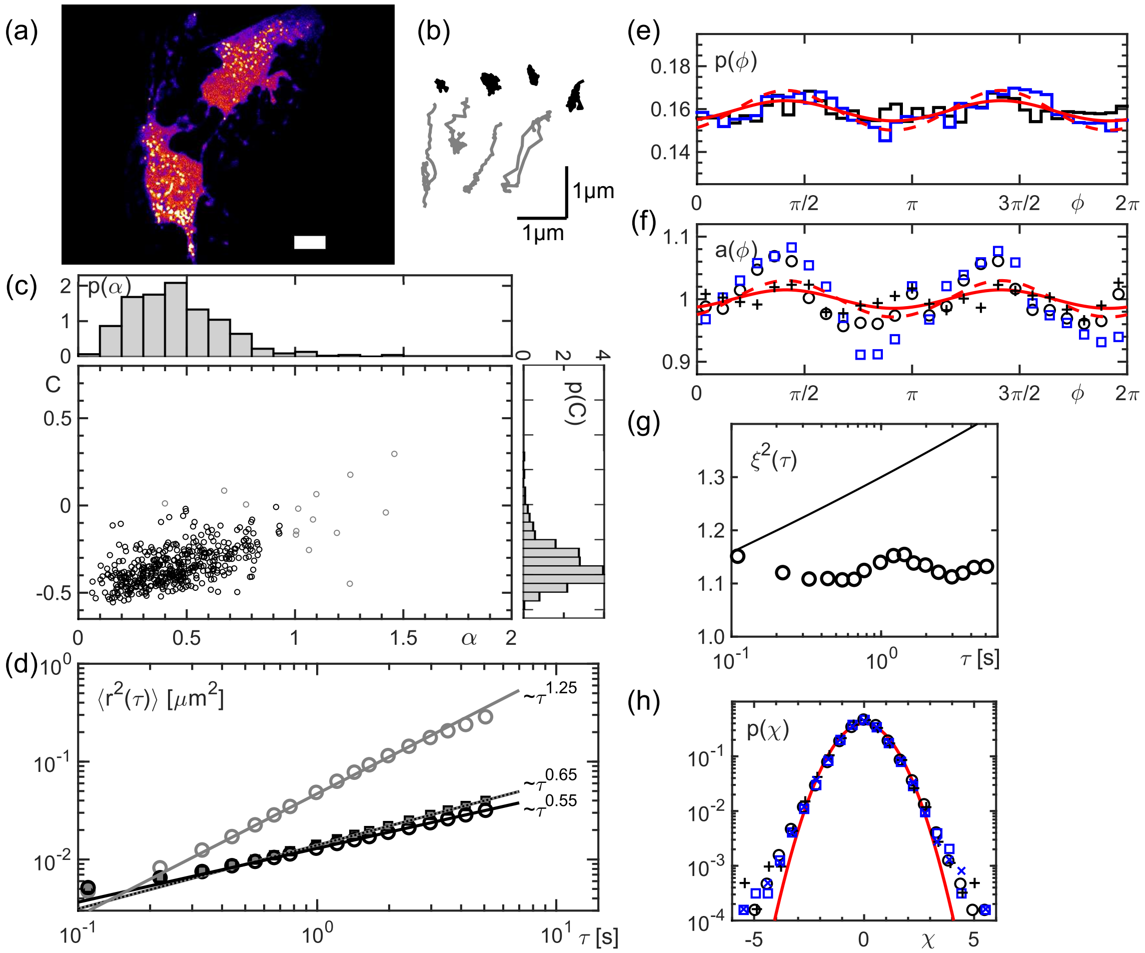


FIG. S2 (a-d) and (e-h) show the analog to Figs. 1 and 2 (main text) and Fig. S1 for a cell in which actin filaments were disrupted by latrunculin A; see also the associated supplementary movie. (c,d) As compared to untreated cells, a reduction of super-diffusive trajectories and a considerable decrease of the scaling exponents for the EA-TA-MSDs of the entire peroxisome ensemble and the sub-diffusive sub-ensemble are observed. (e,f) The modulation in $p(\phi)$ for this particular cell is rather weak, although the ensemble of latrunculin-treated cells still shows a clear modulation on average (cf. Fig. 3 in the main text). Surprisingly, the modulation in $a(\phi)$ is much more pronounced. This somewhat counterintuitive finding is explained by a small fraction of fairly large anisotropic steps that dominate the arithmetic average. Using the median instead (black crosses, to be contrasted to black circles), the over-representation of these steps is avoided and a weak modulation, consistent with that of $p(\phi)$, is regained. (g,h) Similar to nocodazole-treated cells (Fig. S3), the squared step size anistropy, $\xi^2(\tau)$ stays roughly constant (full line from Fig. 2 shown for comparison), and the PDF of normalized step increments, $p(\chi)$, also has the same features as untreated cells (cf. Fig. 2 of the main text).

**Fig. S3**

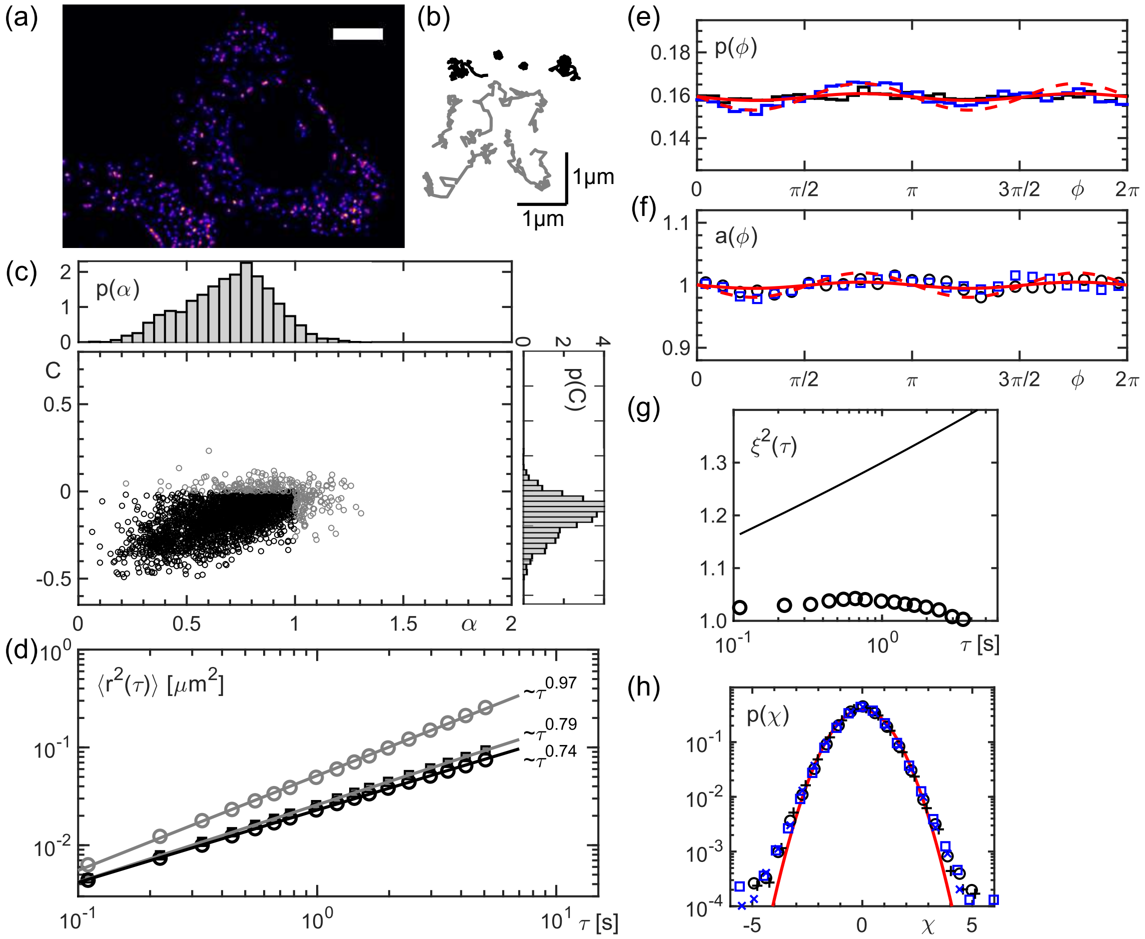


FIG. S3 (a-d) and (e-h) show the analog to Figs. 1 and 2 (main text) and Fig. S1 for a cell in which microtubules were disrupted by nocodazole; see also the associated supplementary movie. (c,d) As compared to untreated cells, a marked reduction of super-diffusive trajectories and a decrease of the scaling exponents for the EA-TA-MSDs of the entire peroxisome ensemble and the super-diffusive sub-ensemble are seen. (e,f) The modulation in $p(\phi)$ and in $a(\phi)$ for this particular cell is strongly reduced in comparison to the shown example of an untreated cell. Still, the ensemble of nocodazole-treated cells shows a clear modulation in both quantities on average (cf. Fig. 3 in the main text). (g) Similar to latrunculin-treated cells, the squared step size anistropy, $\xi^2(\tau)$ stays roughly constant and does not show an increase like untreated cells (full line from Fig. 2 shown for comparison). (h) The PDF of normalized step increments, $p(\chi)$, for steps $\parallel \mathbf{e}_{\phi_0}$ and $\perp \mathbf{e}_{\phi_0}$ is in agreement with the findings for untreated cells, also featuring marked deviations from a normal distribution (red line) in all cases (color-code as in Fig. 2 of the main text).

**Fig. S4**

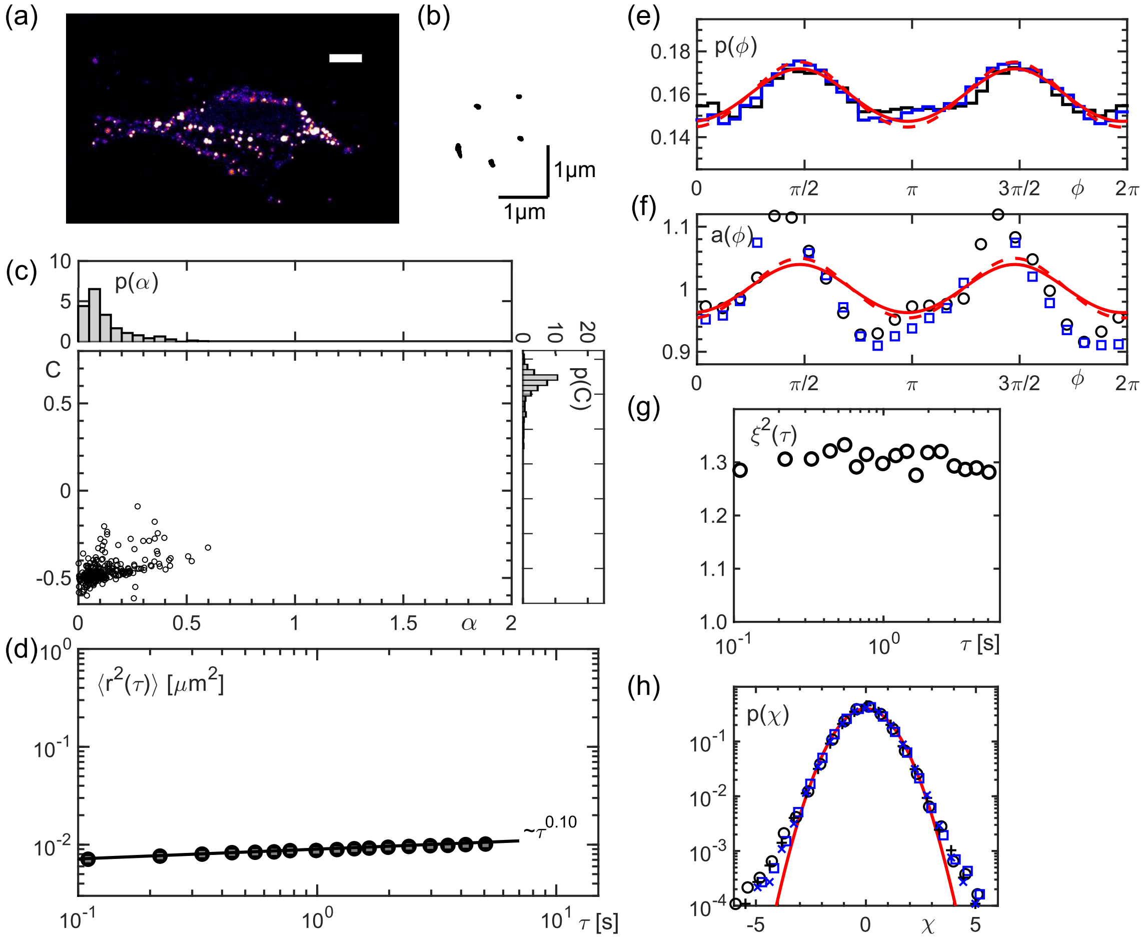


FIG. S4 (a-d) and (e-h) show the analog to Figs. 1 and 2 (main text) and Fig. S1 for a cell in which microtubules and actin were disrupted by treatment with nocodazole and latrunculin; see also the associated supplementary movie. (c,d) As compared to cells that have been left untreated or in which only one filament class was affected, the motion of peroxisomes was almost stalled, yielding only strongly subdiffusive trajectories. (e,f) Despite the heavily reduced step length, the modulation in $p(\phi)$ and in $a(\phi)$ for this particular cell was similar to an untreated cell. (g) In contrast to untreated cells, the squared step size anisotropy, $\xi^2(\tau)$ stays roughly constant and does not show an increase like untreated cells. (h) The PDF of normalized step increments, $p(\chi)$, for steps $\parallel \mathbf{e}_{\phi_0}$ and $\perp \mathbf{e}_{\phi_0}$ is in agreement with the findings for untreated cells, also featuring marked deviations from a normal distribution (red line) in all cases (color-code as in Fig. 2 of the main text).

**Fig. S5**

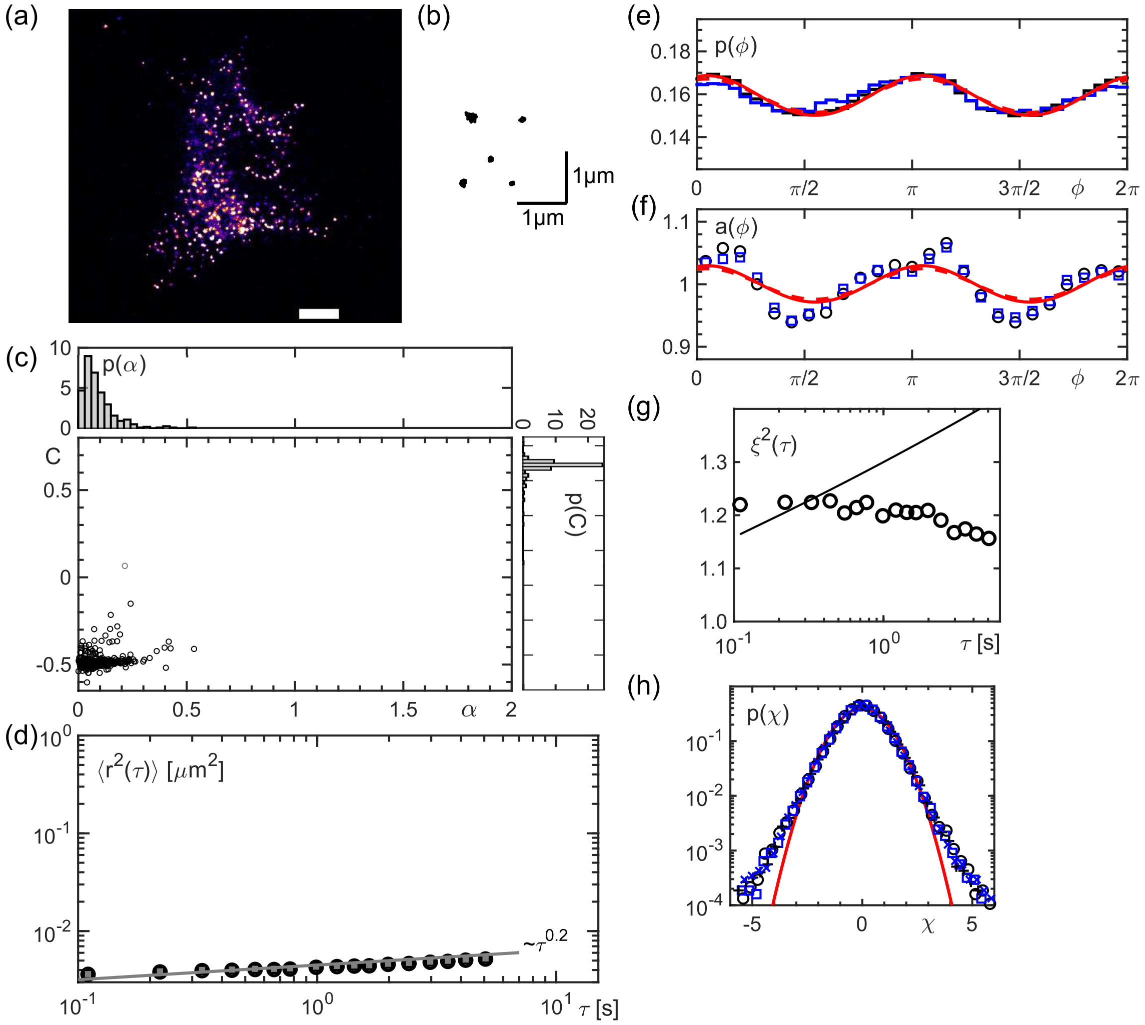


FIG. S5 (a-d) and (e-h) show the analog to Figs. 1 and 2 (main text) and Fig. S1 for a cell in which the endoplasmic reticulum has been scatterd by treatment with filipin; see also the associated supplementary movie. (c,d) Similar to observations in cells that have been treated with both, latrunculin and nocodazle, the motion of peroxisomes was almost stalled, yielding only strongly subdiffusive trajectories. (e,f) Despite the heavily reduced step length, the modulation in $p(\phi)$ and in $a(\phi)$ for this particular cell was similar to an untreated cell. (g) In contrast to untreated cells, the squared step size anisotropy, $\xi^2(\tau)$ stays roughly constant and does not show an increase like untreated cells (full line from Fig. 2 shown for comparison). (h) The PDF of normalized step increments, $p(\chi)$, for steps $\parallel \mathbf{e}_{\phi_0}$ and $\perp \mathbf{e}_{\phi_0}$ is in agreement with the findings for untreated cells, also featuring marked deviations from a normal distribution (red line) in all cases (color-code as in Fig. 2 of the main text).

**Fig. S6**

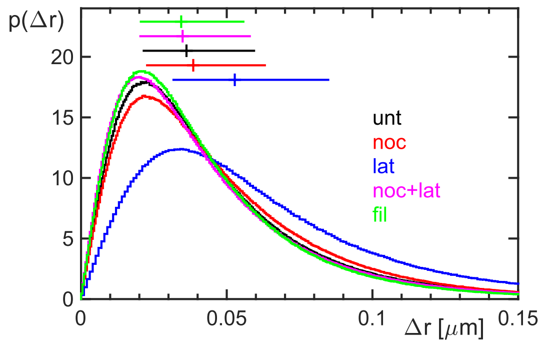


FIG. S6 The PDF of step increment lengths, $p(\Delta r)$ is fairly broad for all conditions (in accordance with the non-Gaussian shape of $p(\chi)$ in Figs. 2d, S2h, S3h, S4h, and S5h). While data for untreated and nocodazole-treated cells are virtually indistinguishable, step lengths are markedly increased in the absence of actin filaments (blue histogram). Combined treatment with nocodazole and latrunculin (magenta) or treatment with filipin yield slightly smaller step lengths as compared to untreated cells. Crosses above the PDFs indicate the respective mean and standard deviation.

**Fig. S7**

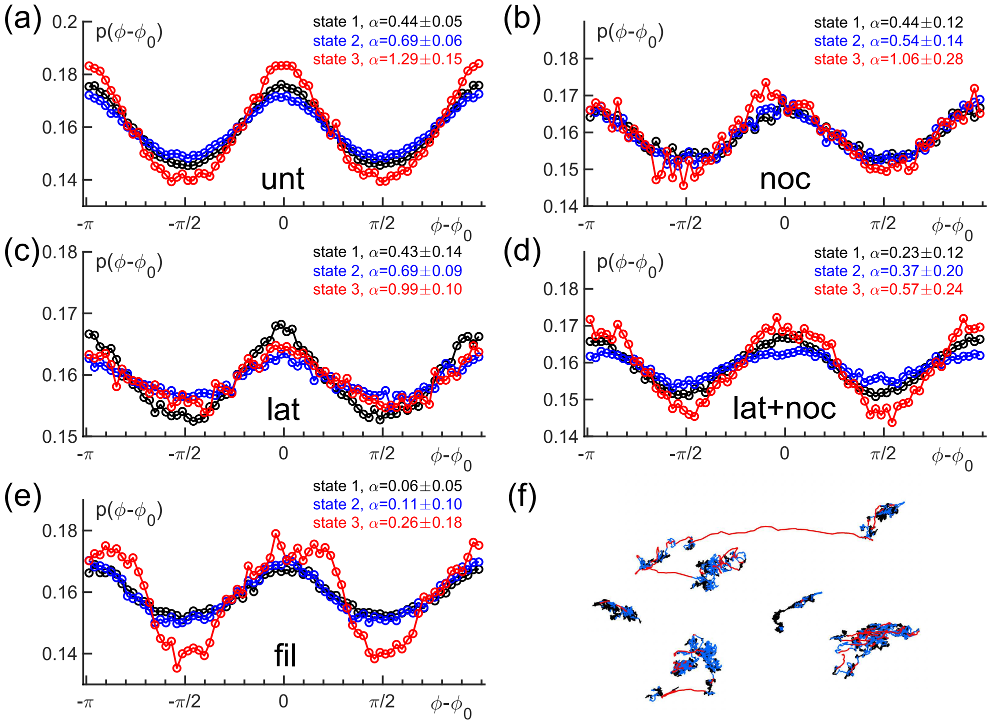


FIG. S7 The PDFs $p(\phi - \phi_0)$, obtained by a HMM model that separates steps into three distinct states (see description above), show for all states and all treatment conditions a clear modulation, hence highlighting a significant anisotropy in the motion. Especially state 2, which by design was meant to attract all steps with a simple and isotropic diffusion, shows the same modulation as the other two states in all cases. Abbreviations in the subfigures (a)-(e) indicate the respective condition (unt: untreated; noc: treatment with nocodazole; lat: treatment with latrunculin A; lat+noc: treatment with nocodazole and latrunculin A; fil: treatment with filipin). (f) Representative trajectories with color-coded segments according to the three states in the HMM classification.
